# Structure and host specificity of *Staphylococcus epidermidis* bacteriophage Andhra

**DOI:** 10.1101/2022.07.21.500982

**Authors:** N’Toia C. Hawkins, James L. Kizziah, Asma Hatoum-Aslan, Terje Dokland

## Abstract

*Staphylococcus epidermidis* is an opportunistic pathogen of the human skin, often associated with infections of implanted medical devices. An increase in antibiotic resistance in *S. epidermidis* and other bacterial pathogens has led to renewed interest in the use of bacteriophages as an alternative to conventional antibiotics. Staphylococcal picoviruses are a group of strictly lytic, short-tailed bacteriophages with compact genomes that are attractive candidates for therapeutic use. Here, we report the structure of the complete virion of *S. epidermidis*-infecting phage Andhra, determined using high-resolution cryo-electron microscopy, allowing atomic modeling of the capsid and tail proteins, including twelve trimers of a unique receptor binding protein, the hexameric tail knob that acts as a gatekeeper for DNA ejection, and the tail tip, which is a heterooctamer of two different lytic proteins. Our findings elucidate critical features that enable host recognition and penetration, facilitating the development of this group of phages for therapeutic applications.

## Introduction

Staphylococci are dominant constituents of the skin microbiome that play critical roles in health and disease. Of the more than 40 skin-associated *Staphylococcus* species (*1*), *S. aureus* and *S. epidermidis* have the greatest pathogenic potential. *S. aureus* is a major cause of a wide range of diseases, from skin and soft tissue infections to lethal sepsis and bacteremia (*2–5*). Although *S. epidermidis* is largely regarded as a beneficial commensal organism, it is also a leading cause of infections of indwelling medical devices (*6–10*).

The emergence of antibiotic resistant strains of *S. aureus* (*11*) and *S. epidermidis* (*9, 12, 13*), combined with a paucity of new antibiotics in the development pipeline (*14–16*), have led to a renewed interest in the therapeutic use of bacterial viruses, or bacteriophages, to eradicate staphylococcal infections (*17–21*). However, many biological challenges, safety concerns, and regulatory hurdles impede the routine use of phage therapy in the US and Europe (*19, 22–24*). Phages often infect a narrow range of hosts, and bacteria can readily evolve resistance to phages, either through the passive acquisition of random mutations in genes essential for phage infection, or by acquiring active mechanisms of anti-phage defense (*22*). To surmount these issues, mixtures of phages with different mechanisms of action are often employed as therapeutic cocktails. Although more effective, such cocktails exacerbate safety concerns pertaining to the preponderance of uncharacterized phage-encoded genes that may cause undesirable side effects. Another concern is the possibility of transduction and mobilization of mobile genetic elements carrying virulence factor genes. These issues underscore the importance of concerted efforts to characterize the mechanisms and impacts of phages, especially those employed as therapeutics.

The picoviruses are a subgroup of the *Podoviridae* family (podoviruses) of short-tailed phages (*25*) that includes the recently designated families *Rountreeviridae* and *Salasmaviridae* (*26, 27*). The most well described member of this group is *Bacillus* phage ϕ29 (family *Salasmaviridae*) (*28, 29*), but several members that infect *S. aureus* are also known, including 44AHJD and P68 (family *Rountreeviridae*) (*30*). Picoviruses are strictly lytic, with genomes of only ≈18-20 kbp (Fig. 1), and employ a unique replication and genome packaging strategy based on monomeric genomes with terminal proteins covalently bound to their 5’ ends (*31*). These features make picoviruses attractive for therapeutic applications. However, picoviruses are underrepresented in phage collections and thus have remained relatively understudied and underutilized.

**Figure 1.**
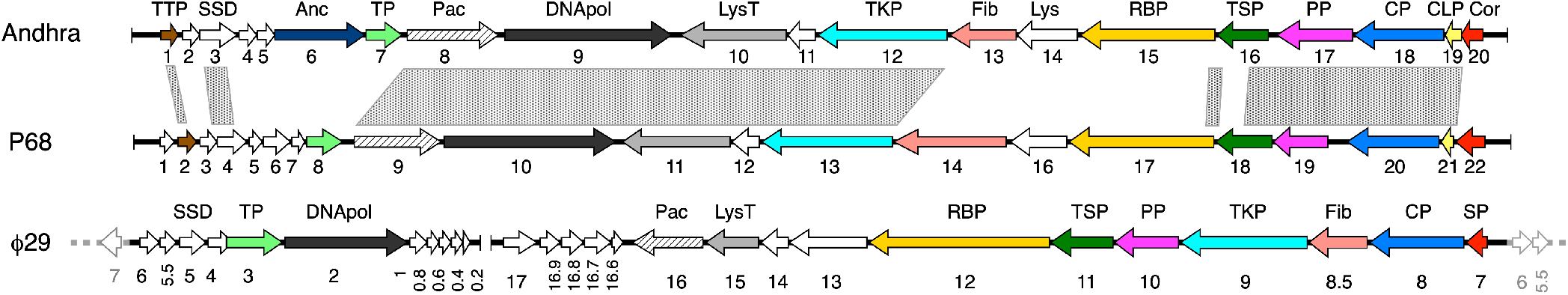
Comparison of Andhra, P68 and ϕ29 genomes. ORFs are numbered in each genome, and relevant genes are labeled. Genes encoding structural proteins are color coded: Tail tip protein (TTP), brown; RBP anchor (Anc), dark blue; terminal protein (TP), light green; tail tip lysin (LysT), gray; tail knob protein (TKP), cyan; head fiber (Fib), salmon; RBP, gold; tail stem/collar protein (TSP), green; portal protein (PP), magenta; major capsid protein (CP), blue; capsid lining protein (CLP), yellow; core (Cor)/scaffolding (SP) protein, red. Non-structural proteins include the packaging ATPase (Pac), hatched; the DNA polymerase (DNApol), black; and the single-stranded DNA binding protein (SSD) and endolysin (Lys), white. The stippled areas between the Andhra and P68 genomes indicate regions of >70% nucleotide identity, calculated in BLASTN (*32*). The ϕ29 genome was split between ORFs 7 and 6, and the right end was moved to the left side (indicated by the duplicated ORFs in gray) to facilitate a more direct comparison to Andhra and P68.

We recently described several new staphylococcal picoviruses, including the *S. epidermidis*-infecting phage Andhra (*32, 33*). These phages are related to other staphylococcal picoviruses in family *Rountreeviridae* (*26*) and share a similar genome organization, with high conservation of core genes involved in DNA replication, packaging and virion structure (Fig. 1). However, a comparison of the genomes of phages Andhra and P68 revealed regions of divergence in Andhra open reading frames (ORFs) 2–7 and ORFs 13–15 (*32*), which likely reflect host-specific differences.

A major determinant of phage host specificity are the bacterial surface structures that serve as phage receptors. In the case of staphylococci and other Gram-positive bacteria, the primary receptor is often wall teichoic acid (WTA), a variable polymer present on the surface of most Gram-positive cells (*34, 35*). Phages bind to these surface structures using a variety of fibers or receptor binding proteins (RBPs) present on their virions. However, the structural basis for host specificity between phages that infect *S. aureus* and *S. epidermidis* is still poorly understood.

Here, we have determined the high-resolution structure of the *S. epidermidis*-infecting phage Andhra virion using cryo-electron microscopy (cryo-EM), allowing atomic models to be built for the capsid and tail structural proteins, including the RBPs and others involved in penetration of the host cell wall. Comparison to P68 allows host-related differences to be examined. Andhra exhibits several unique features compared to P68, including a distinct RBP and a unique lytic protein that also serves to anchor the RBPs to the tail. Unlike previous picovirus structures, we were able to resolve the knob protein hexamer, which acts as a gatekeeper for DNA ejection, and the tail tip, which is formed by a heterooctamer of two proteins. Our data indicate that the structure of the RBPs reflect the type of WTA of the host, and suggest a mechanism for cell wall penetration. A better understanding of host range and the infection process will facilitate the development of this group of phages for therapeutic applications.

## Results

### Overall structure and protein composition of the Andhra virion

Phage Andhra was produced by infection of *S. epidermidis* strain RP62a and purified on CsCl gradients. Phage proteins were separated by SDS-PAGE, treated with trypsin or chymotrypsin, analyzed by mass spectrometry, and compared to the predicted ORFs from the Andhra genomic sequence (Fig. 1). Gene products corresponding to 19 out of the 20 ORFs were identified in the sample, including several expected capsid and tail proteins (Table 1). Some of the detected proteins, such as the DNA polymerase and the packaging ATPase, are not structural proteins *per se*, but likely co-purified with the virions through association with the phage DNA. Proteins encoded by ORF1 and ORF11 were only observed with chymotrypsin cleavage; otherwise the results were similar with both proteases. Several host proteins were also detected, and assumed to be contaminants (not shown).

**Table 1.** Proteins of phage Andhra.

| Andhra ORF | P68 ORF | Amino acids | Seq. Id. (%)<sup>*</sup> | MS Count<sup>†</sup> |  | Oligomer ‡ | Copies/virion | Short name | Function and Location<sup>§</sup> |
| --- | --- | --- | --- | --- | --- | --- | --- | --- | --- |
|  |  |  |  | Tryp | Chym |  |  |  |  |
| 1 | 2 | 90# | 61.1 | N.D. | 3 | 2 | 6 | TTP | Tail Tip Protein |
| 2 | 3 | 71 | 32.4 | 2 | 3 | – | – | – | – |
| 3 | 4 | 169 | 52.4 | 4 | 6 | – | – | SSD | ssDNA Binding Protein |
| 4 | – | 69 | – | 2 | 4 | – | – | – | – |
| 5 | – | 81 | – | N.D. | N.D. | – | – | – | – |
| 6 | – | 403# | – | 61 | 62 | 2 | 12 | Anc | RBP Anchor; Hydrolase |
| 7 | 8 | 167 | 25.2 | 7 | 7 | 1 | 2 | TP | Terminal Protein |
| 8 | 9 | 419 | 65.5 | 39 | 39 | – | – | Pac | Encapsidation Protein; Packaging ATPase |
| 9 | 10 | 763 | 66.2 | 16 | 22 | – | – | DNApol | DNA Polymerase |
| 10 | 11 | 473 | 68.3 | 18 | 19 | 2 | 2 | LysT | Tail Tip Lysin (Spike) |
| 11 | 12 | 136 | 64.7 | N.D. | 5 | – | – | Hol | Holin |
| 12 | 13 | 588 | 69.4 | 138 | 146 | 6 | 6 | TKP | Tail Knob Protein |
| 13 | 14 | 285 | 32.3 | 99 | 99 | 3 | 15 | Fib | Head Fiber |
| 14 | 16 | 299 | 20.1 | 2 | 3 | – | – | Lys | Endolysin |
| 15 | 17 | 609 | 23.5 | 166 | 166 | 3 | 36 | RBP | Receptor Binding Protein; Appendages (Tail Fiber) |
| 16 | 18 | 278# | 44.0 | 39 | 42 | 12 | 12 | TSP | Tail Stem Protein (Lower Collar) |
| 17 | 19 | 335# | 62.1 | 49 | 53 | 12 | 12 | PP | Portal Protein (Upper Collar) |
| 18 | 20 | 405 | 75.3 | 874 | 880 | 3/5/6<sup>+</sup> | 235 | CP | Major Capsid Protein |
| 19 | 21 | 63 | 60.0 | 8 | 8 | 1 | 235 | CLP | Capsid Lining Protein (Inner Capsid) |
| 20 | 22 | 107 | 34.0 | 5 | 5 | 3/6<sup>+</sup> | 72 | Cor | Portal-Proximal Core Protein (Inner Core) |

<sup>\*</sup>Sequence identity between equivalent proteins in Andhra and P68. Calculated using Clustal Omega. A dash (–) indicates that there is no obvious equivalent in P68.
<sup>†</sup>Normalized spectrum counts measured after either trypsin (Tryp) or chymotrypsin (Chym) cleavage. N.D.=not detected.
<sup>‡</sup>In cases where the oligomeric state is ambiguous (e.g. CP), several numbers are given.
<sup>§</sup>Where the names of the equivalent proteins in Andhra and P68 diverge, the P68 name is given in parentheses.

Andhra virions were prepared for cryo-EM by standard methods (Fig. S1A). A data set of 8,818 images was collected on an FEI Titan Krios electron microscope with a Gatan K3 detector, yielding a total of 230,714 particle images (Table 2). These data were subjected to 3D reconstruction in RELION-3 (*36*) without any symmetry imposed (C1). 3D classification revealed that the sixfold symmetric tail existed in two orientations relative to the head, rotated by 30° to each other. Presumably this ambiguity was due to interaction of the sixfold tail with the twelvefold symmetric portal. The final asymmetric reconstruction of the virion therefore used only a subset of 62,061 particle images, and reached 4.43 Å resolution by the Fourier shell correlation (FSC) = 0.143 criterion, according to “gold standard” methods (Fig. S1B).

**Table 2.**
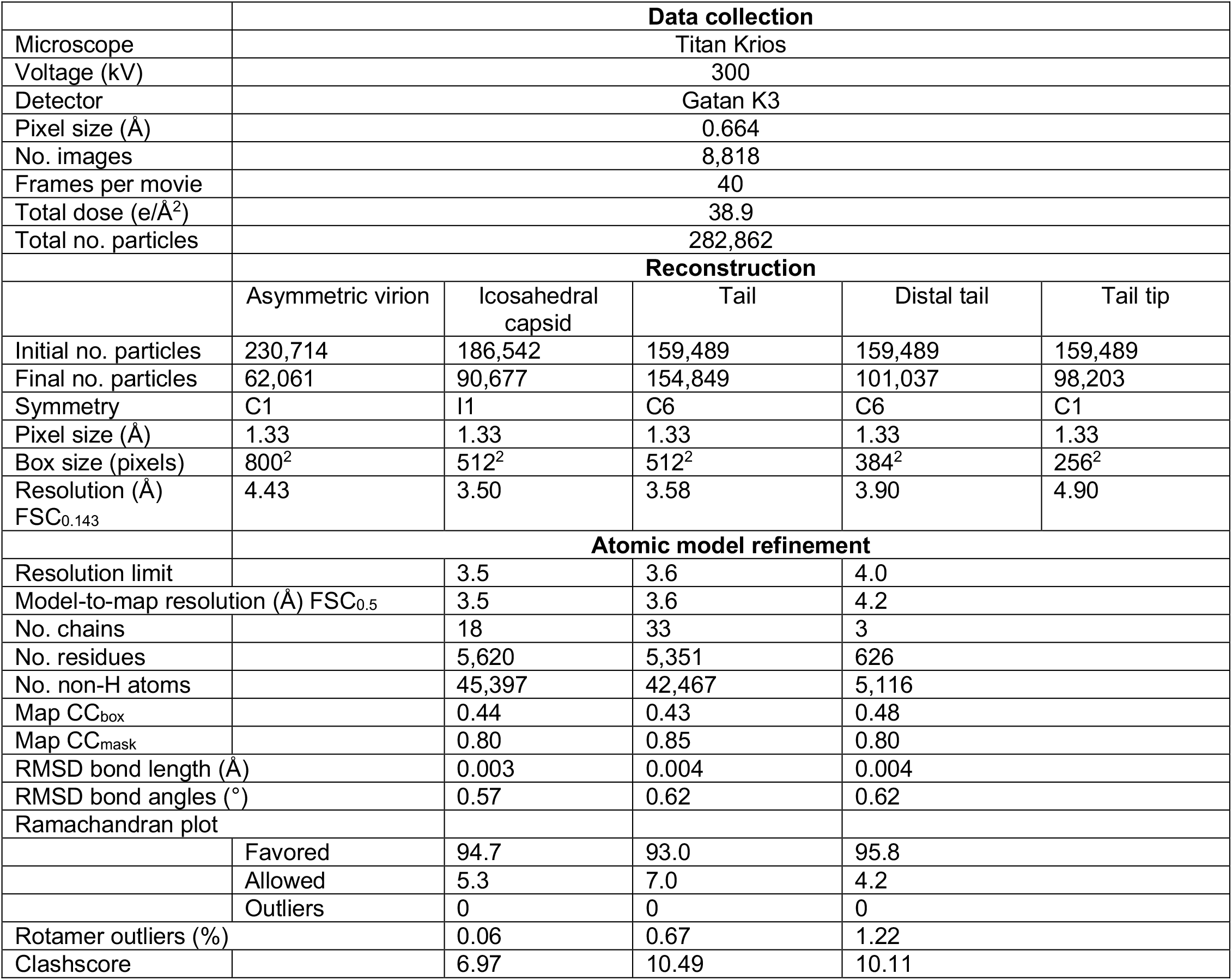
Data collection, reconstruction and refinement data.

The overall structure of the Andhra virion follows that of other phages in this group, with an isometric head (capsid) around 500 Å in diameter (vertex to vertex), connected to a 400 Å long, straight tail surrounded by twelve appendages (Fig. 2A). The head has icosahedral symmetry, with the tail attached to a portal structure located at one of the twelve fivefold vertices. Five trimeric head fibers are attached to the head around the portal vertex. The tail consists of a ≈150 Å wide collar closest to the portal, followed by a ≈160 Å long, 55 Å wide, straight rod that widens into a 145 Å long, 105 Å wide knob (Fig. 2A). The twelve appendages are organized around the rod in a staggered pattern with sixfold rotational (C6) symmetry (Fig. 2B). The asymmetric reconstruction did not resolve an additional bulbous tail tip, whose structure was subsequently determined through focused reconstruction (see below).

**Figure 2.**
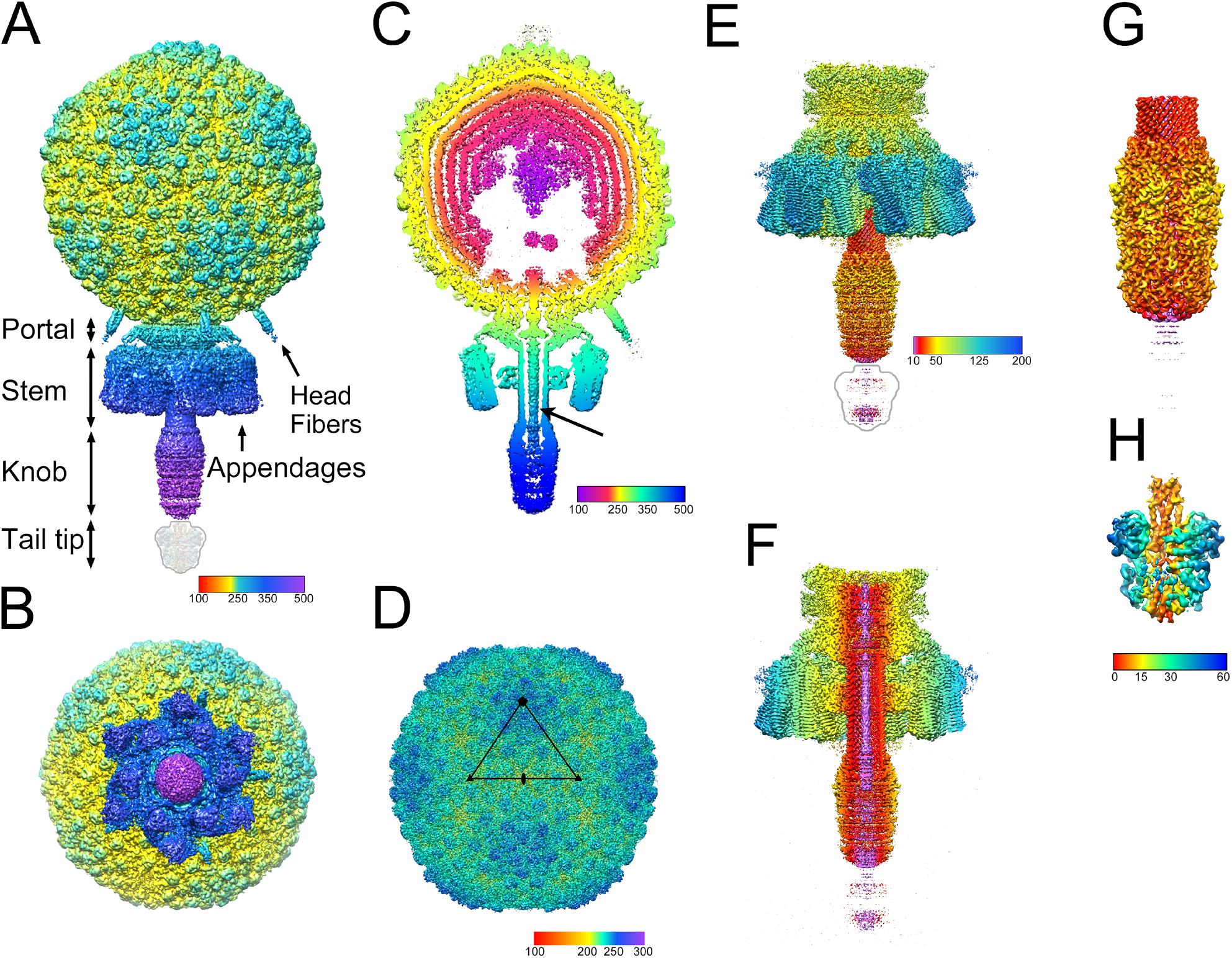
Isosurface representations of the reconstructions. **(A)** Asymmetric reconstruction of the whole virion, colored radially from the center of the capsid according to the color bar (distances in Å). Rendered at 5 standard deviations above the mean (5σ). The tail tip was not visible in the reconstruction at this cutoff, and is indicated by the outline. **(B)** The asymmetric reconstruction viewed from the bottom (tail end), showing the sixfold symmetric arrangement of the appendages. **(C)** 50 Å thick slab through the asymmetric reconstruction, showing the concentric rings of DNA in the interior and the DNA inside the tail (arrow). Colored radially from the center of the capsid according to the color bar. **(D)** Icosahedral reconstruction of the capsid, rendered at 3σ and colored radially from the capsid center according to the color bar. The asymmetric unit triangle is shown, with fivefold, threefold and twofold symmetry axes denoted by the pentagon, triangles and oval, respectively. **(E)** C6 reconstruction of the tail, rendered at 3σ and colored radially from the central axis according to the color bar. The tail tip is indicated by the outline. **(F)** Cutaway view of the tail, rendered and colored as in (E). **(G)** Focused C6 reconstruction of the distal tail, showing the tail knob, rendered at 12σ and colored as in (E). **(H)** Focused asymmetric reconstruction of the tail tip, rendered at 6σ, and colored radially from the central axis according to the color bar.

In cross-section, at least five concentric layers of DNA, spaced about 20 Å apart, are apparent inside the capsid (Fig. 2C). Towards the center of the capsid the layers disappear and are replaced by a more disordered cone-shaped density. Immediately above the portal vertex there is a void, reflecting a lack of consistently organized DNA in this area (Fig. 2C). A 20 Å wide, rod-shaped density, presumably corresponding to the double-stranded DNA and the covalently attached terminal protein, extends from the capsid interior, through the portal and tail and partly into the knob.

### The capsid structure

Reconstruction of the Andhra capsid with the application of icosahedral symmetry using 90,677 particles reached a resolution of 3.50 Å (Table 2; Fig. S1B; Fig. 2D), sufficient to allow atomic modeling. The capsid has *T*=4 architecture and is comprised of 240 copies of the major capsid protein (CP), gene product (gp) of ORF18 (gp18), arranged into 12 pentamers located on the fivefold axes and 20 hexamers located on the twofold axes (Fig. 2D, 3A,B). (In the virion, one pentamer is replaced by the dodecameric portal at the unique vertex.) In addition, there are 240 copies of a small 63-residue protein, gp19, that we refer to as the “capsid lining protein” (CLP). There are thus four copies of CP in the icosahedral asymmetric unit, labeled A–D, with A forming the pentamers and B–D forming the hexamers, and four copies of CLP, labeled E–H (Fig. 3B).

**Figure 3.**
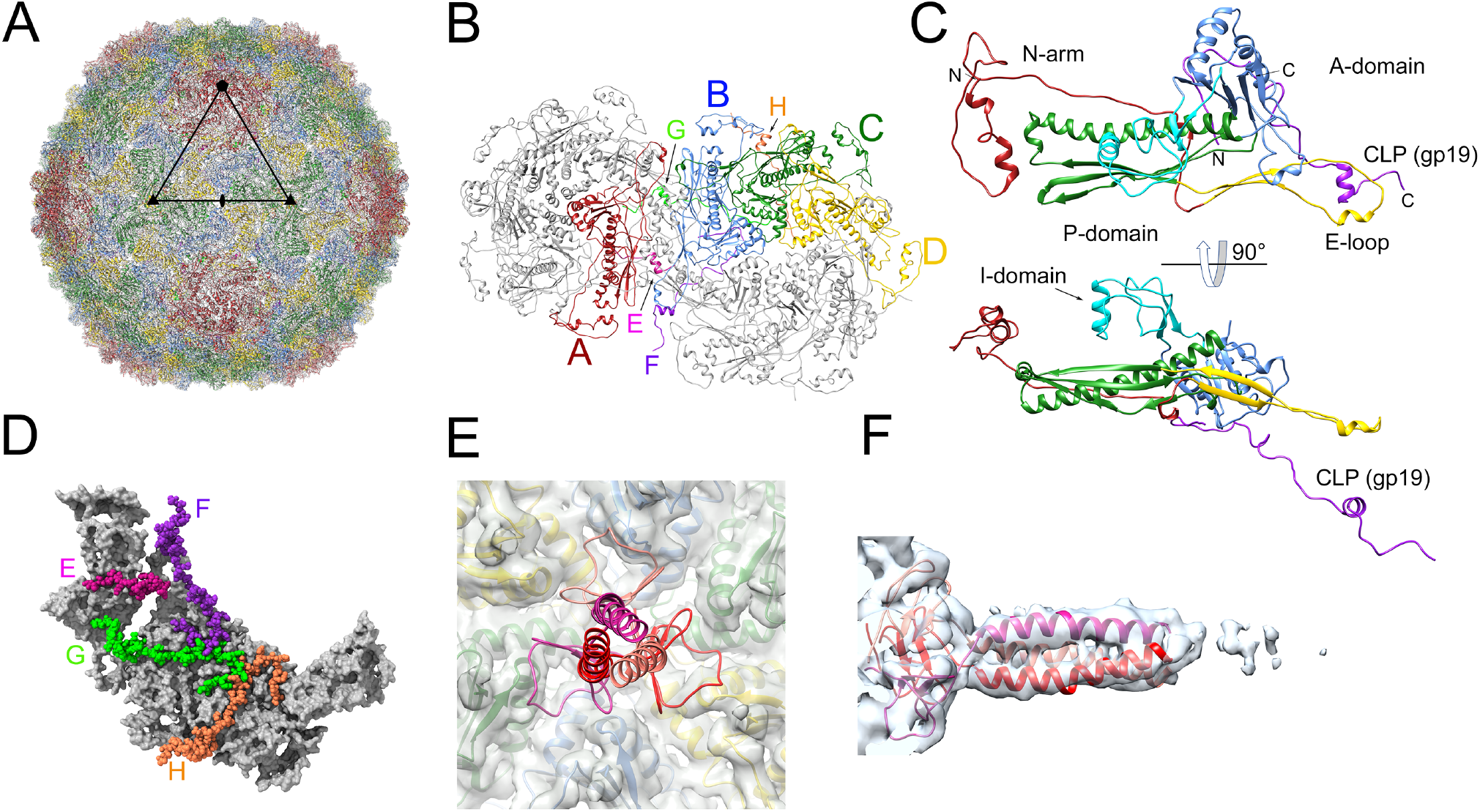
The capsid. **(A)** Ribbon diagram of the entire icosahedral capsid, viewed down a twofold axis, as in Fig. 2D. CP subunits are color coded: A, red; B, blue; C, green; D, yellow. An asymmetric unit triangle, delimited by a fivefold, a twofold and two threefold axes is shown. **(B)** Ribbon diagram showing one hexamer and one pentamer, viewed from the outside of the capsid. The four CP subunits in one asymmetric unit are colored as in (A). The corresponding CLP (gp19) subunits are color coded: E, pink; F, purple; G, light green; H, orange. **(C)** Ribbon diagram of CP subunit B, colored by domain: N-arm, red; E-loop, yellow; P-domain, green; A-domain, blue; insertion domain, cyan. The corresponding CLP (subunit F) is purple. The bottom panel is rotated 90°. **(D)** One asymmetric unit, viewed from the inside of the capsid, showing the CLPs, colored as in (B) with atoms represented as spheres. CP is shown as a gray molecular surface. **(E)** Ribbon diagram of the head fiber (gp13) trimer model (scarlet, ruby and crimson), showing its attachment to the CP hexamer. Capsid density is shown as a transparent gray isosurface, with the corresponding CP subunit models inside. **(F)** Ribbon diagram of the head fiber trimer fitted into the asymmetric reconstruction density (transparent surface), viewed perpendicularly to the axis of the fiber.

The Andhra CP (405 residues; Table 1) has the expected HK97-like fold found in all *Caudovirales*, consisting of an N-arm (residues 1–82), an E-loop (83-122), a P-domain (123-180 and 344-386) featuring a long “spine” α-helix, and an A-domain (181-343 and 387-395) (Fig. 3C). The CP A-domains form tight pentameric and hexameric clusters around the fivefold and twofold (quasi-sixfold) axes, respectively (Fig. 3A, B; Fig. S2). The N-terminal 50 residues constitute a folded sub-domain consisting of two short α-helices and a loop (Fig. 3C). This domain interacts with the equivalent domain from a neighboring CP subunit in a quasi-twofold arrangement (Fig. S2B). There is a 62-residue insertion (I-domain) into the A-domain (residues 276-328) that forms 240 prominent protrusions on the surface of the shell (Fig. 2D, 3A,C). The I-domains are engaged in intra-capsomeric contacts with the E-loop and the N-arm of two threefold related subunits (Fig. S2C). At the icosahedral threefold axes, the P-domains of three C subunits together with an α-helix contributed by three D subunit E-loops come together in a tight ring (Fig. S2D). At the quasi-threefold axes, the P-domains from the A, B and D subunits, together with the E-loops of the A, B and D subunits form a similar structure (Fig S2E,F).

CLP (gp19) forms an S-shaped loop that lines the inside of the capsid (Fig. 3B,C,D). For subunits F– H, the N-terminal 28 residues form a hook-like structure that interacts with the A-domain of a CP subunit. The chain then passes underneath the neighboring subunit within the hexamer and forms a small α-helix that connects to a CP subunit in the adjacent capsomer (Fig 3D). For subunit E, which starts in the pentamer (CP subunit A), only residues 31-53, which connect with a D subunit in the hexamer, were seen (Fig. 3D). On comparing our model to the equivalent protein (gp21) from P68 (*37*), we noticed that the two proteins ran in opposite directions. We therefore re-examined the deposited P68 density (EMD-4442) and found that when the direction of the protein was switched, it showed a better fit (Fig. S3) with the fraction of atoms outside density decreasing from 30.7% to 18.7% and the model-to-map correlation increasing from 0.61 to 0.74.

In P68, gp21 was proposed to constitute a scaffolding protein (*37*)—proteins involved in capsid assembly that are found in most bacteriophages (*38, 39*). However, based on its structure and location in the capsid, we consider it more likely to be a stabilizing factor, functionally similar to so-called “decoration proteins” in many other systems.

### The head fibers

The asymmetric reconstruction revealed head fibers attached to the five hexamers closest to the portal vertex (Fig. 2A). When rendered at low density cutoff level, fibers appeared at other hexamer positions as well (not shown), suggesting that the fibers might attach in other places, but that there is a strong preference for the portal-proximal positions. The head fibers are not visible in the icosahedral reconstruction (Fig. 2D), both due to their low occupancy (only at 5 out of 30 hexamers) and due to the fact that they are trimers attached at an icosahedral twofold axis, so that the icosahedral symmetry averaging would degrade the density.

The head fibers are trimers of the 285-residue protein gp13, which is considerably shorter in Andhra than the equivalent protein in P68 (gp14; 481 residues), and the two proteins share only 32% sequence identity (Table 1). The visible portion of the head fiber protein consists of a 36-residue N-terminal β-prism domain that interacts with the A-domains of every other CP subunit in the hexamer (Fig. 3E), followed by a 31-residue α-helical coiled coil that extends ≈55Å from the surface of the capsid (Fig. 3F). HHpred analysis (*40*) indicated that the coiled coil continues for another 105 residues, followed by a C-terminal domain that is homologous to receptor binding domains from various phages, including *Listeria* phage PSA (PDB ID: 6R5W) and *Lactococcus* phage TP901-1 (PDB ID: 3D8M). These structures were not visible in the reconstructions of either Andhra or P68, presumably due to high flexibility of the fiber.

### Reconstruction and modeling of the tail

In order to resolve the tail at higher resolution, we employed focused reconstruction using a mask encompassing only the tail structures (including the portal and core proteins). An edited version of the C1 reconstruction of the virion in which the capsid density had been manually erased was used as a starting model. This reconstruction was done with C6 symmetry applied and reached a resolution of 3.58 Å (Fig. S1B, Table 2). In this reconstruction, secondary structure elements and many side chains were clearly resolved in the upper (capsid proximal) part of the tail (Fig. 2E). The lower (distal) part of the tail was not equally well resolved in this map, however, and required separate focused reconstructions (see below).

The C6 reconstruction was used to build atomic models for the proteins in the upper tail, including 72 copies of a small portal-proximal core protein (gp20, 107 residues, referred to as “inner core” in P68), the dodecameric portal protein (PP, gp17, 335 residues; referred to as “upper collar” in P68 and “connector” in ϕ29; see Fig. 1), the dodecameric tail stem or collar protein (gp16, 278 residues; referred to as “lower collar” in P68 and ϕ29), the twelve trimers of RBP (gp15, 609 residues; referred to as “tail fiber” in P68), and six dimers of a protein we identified as the product of ORF6 (gp6, 403 residues; see below) (Fig. 4A–C; Fig. S4).

**Figure 4.**
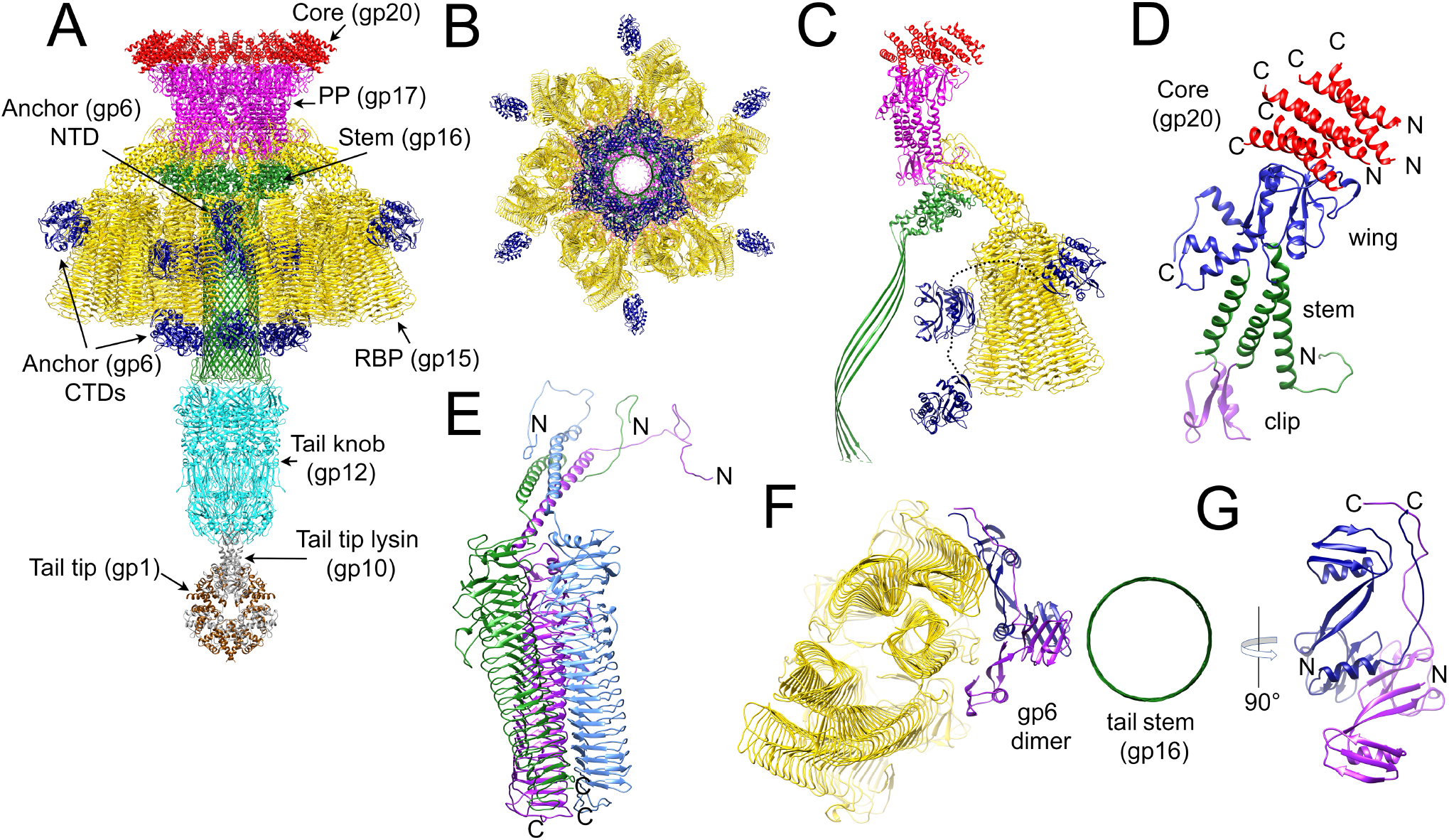
The tail structure. **(A)** Ribbon diagram of the composite model of the Andhra tail proteins, viewed from the side and colored as in Fig. 1: core protein (gp20), red; portal protein (gp17), magenta; tail stem protein (gp16), green; RBP (gp15), gold; RBP anchor (gp6), blue; tail knob protein (gp12), cyan; tail tip lysin (gp10), gray, tail tip protein (gp1), brown. **(B)** Model of the tail, viewed from the bottom; knob and tip not included. **(C)** One asymmetric unit of the C6 tail reconstruction, comprising twelve copies of core protein (gp20), two copies of portal protein (gp17), two copies of tail stem protein (gp16), six copies (two trimers) of RBP (gp15) and two copies of the anchor protein (gp6). (Knob and tip not included.) The connections between the N- and C-terminal domains of gp6 are indicated by the dotted lines. **(D)** Portal protein monomer with the wing, stem and clip domains colored blue, green and purple, respectively. The corresponding 6 copies of the core protein (gp20) are shown in red. N- and C-termini are indicated. **(E)** One RBP trimer, with protomers colored blue, green and purple. The N- and C-termini are indicated. **(F)** Ribbon diagram of a section through the tail, viewed from the top, showing a dimer of gp6 (blue and purple) interacting with the barrel of the tail stem (green) and the RBPs (gold). **(G)** 90° rotated view of the gp6 dimer, viewed perpendicular to the axis of the tail. N- and C-termini are indicated.

### The portal, core and tail stem proteins

The Andhra portal consists of twelve copies of PP (gp17), arranged in a 150 Å wide ring with C12 symmetry, with a 30 Å wide central hole (Fig. 4A,B). There was clear density in the C6 tail reconstruction for PP from residue 6 to 334. The PP monomer consists of three domains: the wing (51–160 and 258–334), the stem (6–50, 161–186 and 234–257) and the clip (residues 187–233) (Fig. 4D). The stem domain consists of three long α-helices that traverse the capsid shell. Residues 6–20 form a loop that interacts with the fivefold symmetric capsid shell, as previously described for P68 (*37*). The wing domain resides inside the capsid where it interacts with the portal-proximal core protein (gp20) and the DNA (Fig. 4C), while the clip domain connects with the tail stem protein (gp16).

The gp20 core protein is associated with the portal on the inside of the capsid (Fig. 4A,C). There was clear density only for residues 42–64, which form a single α-helix. Two bundles of three α-helices sit on top of each copy of the portal protein, for a total of 72 copies of gp20 in the capsid (Fig. 4D). gp20 has only weak homology (34% sequence identity) to the somewhat longer (133 residues) “inner core” protein (gp22) of P68. Based on its location in the genome and overall α-helical characteristic, gp20 might be equivalent to the scaffolding protein (SP; gp7) of ϕ29 (*38, 39, 41*). Unlike a typical scaffolding protein, however, gp20 remains in the mature capsid after DNA packaging.

On the distal side of the portal, the dodecameric tail stem or collar protein, gp16, forms an elongated tube that comprises the main stem of the tail (Fig. 4A,C). We could model gp16 from 38 residues upstream of the annotated ORF16 sequence in the GenBank entry for phage Andhra (KY442063.1), suggesting that the correct start is at nucleotide 15,474 (TTG), adding an extra 39 residues to the sequence, for a total of 278 residues (Table 1). TTG (leucine) is a common alternative start codon in *Staphylococcus*. gp16 consists of a 147-residue globular α-helical collar domain that associates with the clip domain of PP (Fig. 4A,C). A 128-residue insertion between amino acids 122 and 250 forms a twisted β-hairpin (Fig. 4A,C), creating a 150 Å-long tubular β-barrel with an inner diameter of ≈32 Å that acts as a conduit for the DNA. The β-barrel tube is ≈40Å longer in Andhra than in P68, corresponding to an additional 28 residues in the β-hairpin. The tip of the hairpin forms a hook that was not visible in the tail reconstruction, but was modeled in the focused reconstruction of the lower tail (see below).

### The receptor binding proteins

The twelve appendages surrounding the tail (Fig. 2A,E) correspond to trimers of RBP, the 609-residue product of ORF15 (gp15). There are thus 36 copies of RBP in the Andhra tail, arranged in a staggered fashion (Fig. 4A,B). The staggered C6 arrangement is unlike P68, where the twelve RBP trimers are arranged with C12 symmetry (Fig. S4).

The Andhra RBP consists of an N-terminal 36-38-residue loop followed by a 34-38-residue α-helical coiled-coil stem domain, connected to a 534-residue C-terminal β-helix domain (Fig. 4E). The N-terminal loop interacts with the stem and clip domains of PP and with the collar domain of gp16 (Fig. 4C; Fig. S5A,B). The C-terminal β-helix domain, which comprises the majority of the RBP, consists of a stack of 19 triangular layers, each consisting of 2-3 β-strands arranged in a regular, repeating zigzag pattern (Fig. 4E, S5C). At its N-terminal end, of the β-helix is more irregular, with longer insertions between the layers. The β-helix domains of the three RBP subunits in the trimer superimpose with an RMSD=0.4 Å, but the N-terminal loops and coiled-coil domains diverge in order to accommodate the sideways attachment of the RBPs to the tail (Fig. 4E, Fig. S5C). Due to the staggered arrangement of the RBP trimers, there are two types of trimers—proximal and distal—with different organization of the N-terminal domains (Fig. S5D).

While the Andhra RBP is functionally equivalent to the RBP (or “tail fiber”) of P68 (gp17), the proteins share almost no sequence homology and have completely different structures. Whereas P68 gp17 has a structure similar to the RBP of *S. aureus* siphovirus 80α (*42*), the β-helix structure of Andhra RBP is related to alginate lyases and other bacterial enzymes involved in breaking down polysaccharides. Similar proteins are found in several bacteriophages, such as the tail spikes of *Salmonella* phages Det7 (PDB ID 6F7D) (*43*) and P22 (PDB ID: 1TSP) (*44*). Notably, the Andhra RBP is similar to the N-terminal two-thirds of the β-helix domain of the ϕ29 RBP (called “preneck appendage”, gp12; PDB ID: 3GQ9)(*45*). In the reconstructions of P68 and ϕ29, the RBPs were poorly resolved, requiring separate X-ray structures to produce a pseudo-atomic structure for the tail (*37, 46*). In contrast, the entirety of the Andhra RBP structure was well resolved. In spite of the overall distinct structural folds of the Andhra and P68 RBPs, the N-terminal loops are similar (Fig. S5E) and superimpose with an RMSD=1.1 Å over 24 residues, including a stretch of 17 residues that are 100% identical between Andhra and P68 (Fig. S5F). This loop represents the point of interaction between the RBPs and the portal and stem proteins, which are similar in Andhra and P68.

### The RBP anchor protein, gp6

After modeling the Andhra upper tail proteins with equivalents in P68, six well-resolved densities between the RBPs and the tail stem (gp16) remained in the tail reconstruction. Close inspection of the density revealed two tryptophan residues separated by one other residue (a WXW motif) in an αβ-helix. A sequence search revealed that this motif was found in ORF6, encoding gp6, a protein that has no equivalent in P68 (Fig. 1). Using the WXW motif as an anchor point, we were able to build the N-terminal 143 residues of gp6 into the density. Well resolved density extended six residues upstream of the annotated start codon for ORF6 at nucleotide 1,905. The true start for gp6 is thus most likely at nucleotide 1,887 (TTG). gp6 forms six asymmetric dimers that bridge the RBP trimers and the tail stem protein (gp16) (Fig. 4F). The gp6 monomer consists of two α/β subdomains connected via two antiparallel β-strands (Fig. 4G). In the virion, the two monomers stack on top of each other with the C-termini emerging on the same side of the dimer (Fig. 4F,G). This arrangement allows each gp6 dimer to interact with both a proximal and a distal RBP trimer (Fig. 4C,F). Since the gp6 dimers appear to anchor the RBP trimers in place, we refer to this protein as the “RBP anchor” protein. Most likely, stabilization of the RBPs by these anchors was responsible for the well-ordered RBPs in Andhra compared to P68.

The C-terminal ≈200 residues of gp6 are predicted by HHpred to contain a hydrolase domain with homology to bacterial lipases, including the tail-associated lysin (Tal) of bacteriophage 80α (PDB ID: 6V8I) (*42*), suggesting that gp6 is involved in penetration of the cell wall. After building the gp6 N-terminal domains, there was still six unassigned, weak densities attached to the periphery of the RBPs and six densities attached at the distal end of the tail stem. These two types of densities could be superimposed with a correlation of ≈0.9, suggesting that they belonged to the same protein, and the most likely candidate was the C-terminal domain of gp6. Indeed, an AlphaFold (*47*) model for the C-terminal domain of gp6 could be rigid-body fitted into both densities with high correlation (>0.85). The linkers connecting the N- and C-terminal domains could not be resolved, but based on the direction of the C-terminal arms of the N-terminal domains of the two gp6 subunits, the top subunit most likely connects to the peripheral C-terminal domain, while the bottom subunit connects to the distal domain (Fig. 4C).

### The distal tail

It was clear from both the asymmetric reconstruction of the virion and the C6 tail reconstruction that the distal part of the tail was not well resolved (Fig. 2A, E). This part corresponds to the tail “knob” and “tip” (referred to as “spike” in P68). These features were not well resolved in the P68 reconstructions (*37*), and initially appeared to be poorly ordered in Andhra as well. We presumed that this was due to an orientation ambiguity between the distal and upper portions of the tail. Although the tail knob was expected to form a hexamer with C6 symmetry, it was inconsistently aligned relative to the RBPs, with two possible orientations, differing by a 30° rotation. We employed symmetry expansion and 3D classification to generate a masked, focused reconstruction of this portion with only C3 symmetry, which was then re-symmetrized to C6. Refinement of the C6 reconstruction of the distal tail reached a resolution of 3.9 Å, allowing for atomic modeling (Fig. 3G. Table 2).

The tail knob consists of a hollow cylinder, 145 Å long and 110 Å wide, and is made up of a hexamer of gp12 (Fig. 5A,B). gp12 has a complex, intertwined structure that is similar to those of the gp9 tail knob protein of phage ϕ29 (*48*) and the streptococcal phage C1 major tail protein gp12 (*49*), to which it could be aligned with RMSD=1.1 Å for 212 out of 588 residues. The N-terminal part of gp12 forms a -sandwich domain related to tail tube proteins from many members of the *Siphoviridae* and *Myoviridae* bacteriophages, connected to an elongated β-sheet domain at the bottom of the hexamer via an extended connecting loop (Fig. 5C). The polypeptide chain then returns to the top of the hexamer to complete the β-sheet in the N-terminal domain. Residues 387–481 form an extended α-helical loop (the “channel loop”) on the inside of the cylinder, which constricts the inner diameter to ≈12 Å, presumably preventing exit of the DNA and terminal proteins (Fig. 5B,C). This loop was not resolved in the crystal structures of C1 gp12 (*49*) or ϕ29 gp9 (*48*), but corresponding density was observed in the ϕ29 reconstruction (*46*). Most likely, these loops reorganize to become extruded through the tip of the tail during infection, enabling the resulting 12 α-helices to insert into the host membrane to allow the passage of the DNA into the cell, similar to the mechanism proposed for ϕ29 (*46*). The focused reconstruction of the distal tail also resolved the hooks at the end of the β-hairpins of the tail stem protein (gp16) dodecamer that were not seen in the C6 reconstruction of the whole tail (Fig. 5A).

**Figure 5.**
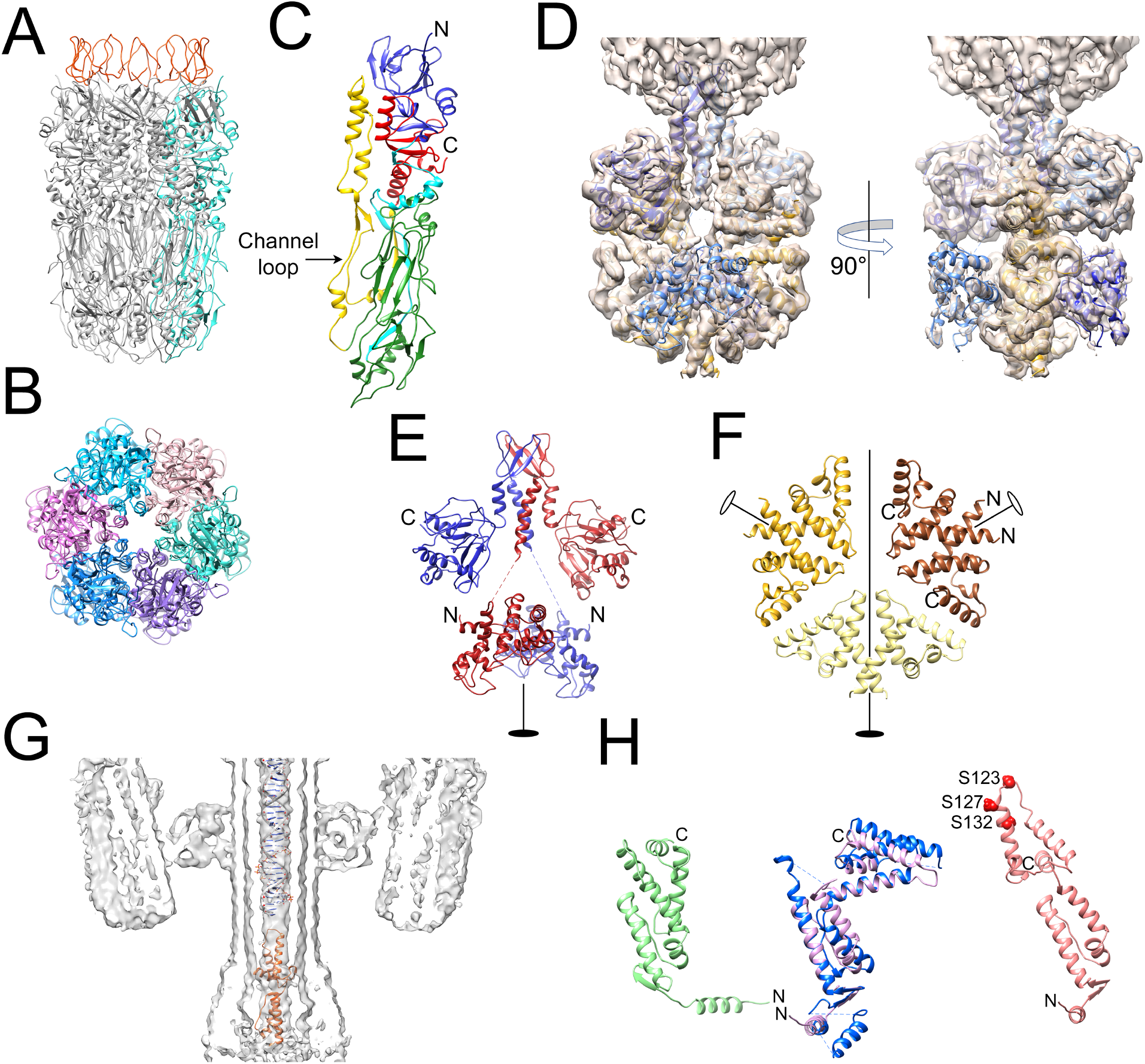
Distal tail structure. **(A)** Ribbon diagram of a hexamer of the tail knob protein (gp12). One protomer is colored cyan. Loops belonging to the tail stem protein, gp16, are shown in red. **(B)** Ribbon diagram of the gp12 hexamer, viewed from the tail tip. Each protomer is colored in a different shade of blue or purple. **(C)** Ribbon diagram of a gp12 monomer, colored by domain: N-terminal domain, blue; N-to-C linker, cyan; tip domain, green; channel loop, yellow; C-terminal domain, red. **(D)** Isosurface of the focused asymmetric reconstruction of the tail tip with the two copies of gp10 and the six copies of gp1 shown in blue and gold, respectively. The right panel is rotated by 90°. **(E)** Ribbon diagram showing the two subunits of gp10 (red and blue). N- and C-termini and the twofold symmetry axis are indicated. **(F)** Ribbon diagram showing the six copies of gp1 in the tail tip arranged as three dimers, colored brown, sand and gold. The twofold symmetry axis is indicated by the filled oval, and the two peripheral dimer twofold axes are shown as open ovals. N- and C-termini are indicated on one dimer. **(G)** 15Å thick slab through the asymmetric virion reconstruction, showing the central density in the tail with double stranded DNA and the gp7 model (orange) fitted in. **(H)** Left panel: Ribbon diagram of the AlphaFold model for gp7 (green). Middle panel: the modified gp7 model (pink) superimposed on the ϕ29 gp3 crystal structure (blue). Right panel: gp7 (salmon) modified to fit into the central tail density. The serine residues near the top of the molecule (S123, S127 and S132) are indicated as red spheres.

The distal tail reconstruction still did not resolve the bulbous density at the tip of the tail (Fig. 2E, G). The tip density was not resolved in the P68 reconstruction either, and was suggested to consist of a pentamer of gp11 (*37*), equivalent to Andhra gp10 (Fig. 1), a protein that we had previously demonstrated to act as a lysin with cell wall hydrolase activity (*32*). To resolve the tip density, we carried out another masked reconstruction focused on the tail tip without the application of symmetry, reaching a resolution of 4.9 Å (Table 2). This reconstruction revealed a twofold symmetric structure consisting mostly of α-helices, with a fourβ-helix bundle inserting into the knob (Fig. 2H; 5D).

A partial crystal structure of the N-terminal domain (residues 2-200) of P68 gp11 was solved previously (PDB ID: 6O43) (*50*), and was shown to be homologous to streptococcal phage C1 lysin PlyCA, for which a full-length structure was previously determined (PDB ID: 4F88) (*51*). The PlyCA structure consists of a glycosyl hydrolase (GyH) domain connected to a Cys-His aminohydrolase/peptidase (CHAP) domain. In phage C1, these enzymatic activities are involved in breaking down the streptococcal cell wall. An AlphaFold model for Andhra gp10 revealed the two expected domains (GyH and CHAP), separated by a helixβ-sheetβ-helix linker domain (residues 211– 277). The three domains were individually fitted into the tail tip reconstruction, revealing a gp10 dimer with the N-terminal GyH domains at the bottom of the tip and the C-terminal CHAP domains at the top (Fig. 5D,E). The two α-helices of the linker domain fit into the connecting densities between the tip and the knob, with the β-sheet penetrating into the gp12 knob protein above (Fig. 5D,E). The density corresponding to the N-terminal domain was weak (Fig. 5D), suggesting low occupancy and high degree of flexibility.

After fitting the gp10 dimer into the tip density, it was clear that the tip contained at least one other, primarily α-helical protein. After subtracting the gp10 density, the remaining density appeared as a flat disc with quasi-threefold symmetry. Closer analysis revealed that the disc was made of three dimers, with the twofold symmetry axis of the tip passing through one dimer and the other two dimers on opposite sides of the tip (Fig. 5D). The shape and size of the density was consistent with a mostly α-helical protein of approximately 90 residues. The only unassigned candidate proteins of the right size were gp1 (90 residues), gp2 (71 residues), gp4 (69 residues) and gp5 (81 residues). We generated AlphaFold models for each of these proteins. gp2, gp4 and gp5 were a poor match for the tail tip density, but gp1 had a compact, α-helical structure that could be fitted as three dimers into the tail tip disc density (Fig. 5D,F). In particular, the C-terminal helix had several aromatic residues that matched bulky side chain densities in the map. We noticed that the predicted N-terminal αβ-helix was too short for the density. Upon re-inspecting the sequence, we realized that the published sequence was missing 11 residues, with the most likely start being a GTG (valine) codon upstream of the annotated sequence in GenBank (KY442063.1). Gp1 has homologues in other staphylococcal picoviruses, including gp2 from P68, but no other homologues that could give an indication of its function, such as an enzymatic activity.

### Internal tail density

On the inside of the tail, a ≈22 Å wide, elongated density stretched from the capsid, through the portal and tail tube to the knob, where it widened slightly (Fig. 2C, F). In the collar at the interface between the portal and tail tube, the density formed a disc-like shape. The central density was not well resolved in either the C6 tail reconstruction or the asymmetric reconstruction of the virion, and could not be modeled. Attempts at masked, focused reconstruction without symmetry were not successful.

In the P68 reconstruction, the internal density was proposed to correspond to a trimer of P68 gp8 (equivalent to gp7 in Andhra), and designated as “needle” based on its high α-helical content and elongated shape (*37*), by analogy with the trimeric needle protein (gp26; PDB ID: 4ZKU) of phage P22 (*52*). However, this similarity is most likely only superficial, since these two systems are highly dissimilar, and the P22 needle is located on the outside of the P22 tail tip.

Instead, the internal density most likely corresponds to one end of the genome. The width of the density matches that of double-stranded DNA (Fig. 5G). The bulbous density inside the knob presumably corresponds to the covalently attached Andhra terminal protein (TP). Based on comparisons with the ϕ29 genome, and excluding ORFs with known functions, the most likely candidate for the Andhra TP is gp7 (Fig. 1). An AlphaFold model of gp7 has an elongated, α-helical structure with the same topology as the ϕ29 terminal protein, gp3, for which a crystal structure bound to the ϕ29 DNA polymerase was previously determined (PDB ID: 2EX3; Fig. 5H) (*53*). The two proteins superimpose with an RMSD of 2.2 Å after rotating the N- and C-terminal subdomains of gp7 (Fig. 5H). Fitting of gp3 into the ϕ29 reconstruction required extensive modification (*46*), suggesting that the protein takes a different, more elongated conformation in the virion compared to the polymerase-bound form. When gp7 is modified similarly (Fig. 5H), it fits roughly into the bulbous density inside the knob (Fig. 5G). In ϕ29, the 5’ end of the DNA is covalently attached to residue S232 of gp3. It is not known which residue is bound to the DNA in Andhra, but in the gp7 model, several serine residues, including S123, S127 and S132, are in a position where they could interact with the DNA (Fig. 5H).

## Discussion

All tailed bacteriophages are considered to belong to the same lineage (the *Caudovirales*), which is evident from similarities in the structures of many key proteins, including the major capsid protein, the portal protein and the major tail proteins (*54*). Traditionally, the *Caudovirales* are divided into three families based on the structure of the tail: long and contractile (*Myoviridae*), long and flexuous (*Siphoviridae*), or short (*Podoviridae*). Picoviruses like Andhra, P68 and ϕ29 have been considered a subfamily (*Picovirinae*) in the *Podoviridae* family due to the presence of a short tail (*25*). However, these phages have many features that distinguish them from other podoviruses, like P22 and T7, including their small genomes and their distinct replication and DNA packaging strategy (*31*). The International Committee on Taxonomy of Viruses recently separated out the picoviruses into three new families (*26, 27*): *Salasmaviridae* (which includes *Bacillus* phage ϕ29 and related viruses like B103 and GA-1 (*29*)), *Guelinviridae*, and *Rountreeviridae* (the latter including Andhra, P68 and 44AHJD (*30*)). While the genomic organization differs somewhat between the *Salasmaviridae* and the *Rountreeviridae*, they are clearly related (Fig. 1). The three-dimensional structures of key proteins are also generally conserved between the two families, with some exceptions where host interactions are involved, e.g. the RBPs. In a fascinating twist to this genealogy, structures of the RBPs of Andhra and P68 show close relationships to phages belonging to divergent lineages: the P68 RBP is highly similar to the RBP of staphylococcal siphovirus 80α (*42*), while the Andhra RBP resembles proteins from phages that infect a variety of hosts, including *Bacillus* (29) and *Salmonella* (Det7). These relationships are a reflection of the extensive horizontal evolution between phages from vastly different lineages.

Assembly of capsids of the *Caudovirales* bacteriophages generally require the assistance of a scaffolding protein, which acts catalytically on the capsid protein the ensure correct assembly (*38, 39*). (One exception are the HK97-like phages, such as *S. aureus* phage 12, in which an N-terminal scaffolding domain of the CP serves the same purpose (*55, 56*).) ϕ29 has a scaffolding protein, gp7, that appears to be similar to those of other phages, including staphylococcal phage 80α and its satellite SaPI1 (*41, 57, 58*); however, its location in the ϕ29 capsid is still uncertain (*46*).

In contrast, Andhra and P68 do not have a direct equivalent to the ϕ29 SP. Structurally, the Andhra protein most similar to a classical SP is the portal-proximal core protein, gp20, which, like other SPs, is predominantly α-helical. The location of ORF20 in the Andhra genome is equivalent to that of the SP gene (ORF7) in ϕ29: at the beginning of the structural operon, preceding the CP gene (Fig. 1). Unlike ϕ29, Andhra and P68 have an additional ORF between that of the core protein and the capsid protein, which was interpreted as the scaffolding protein gene in P68 (*37*). However, this protein—the capsid lining protein—does not resemble a scaffolding protein and more likely serves a stabilizing role. The C-termini of the core proteins form α-helical bundles that interact primarily with the portal (Fig. 4C,D). In P68, the N-terminus of gp21 was shown to interact with CP (*37*), as expected for a scaffolding protein, but we did not observe density for this part in Andhra. There is a precedent for a portal-proximal location of the SP in ϕ29, where gp7 was found to bind to the gp10 portal protein and probably provides a bridge between the portal and the capsid (*59, 60*). Indeed, a key role of scaffolding proteins may be to ensure incorporation of the portal during nucleation of capsid assembly. Unlike a typical scaffolding protein, however, the core protein remains associated with the capsid in the mature virion (*38, 39*). Perhaps the portal-proximal organization we observe in Andhra and P68 is only the subset of the scaffolding that remains in the capsid.

From these observations, it is still uncertain if Andhra and its relatives use a catalytic scaffolding protein during assembly, or if the process is somewhat different from other *Caudovirales* phages. Picoviruses do not undergo the large expansion and major structural transitions associated with capsid maturation and DNA packaging observed in other tailed phages. It is also worth noting that ϕ29 differs from Andhra and P68 in having a prolate, rather than an isometric capsid. Further experiments are needed to ascertain whether the portal-proximal core protein is indeed a scaffolding protein.

Unlike most *Caudovirales* phages, which package concatemeric DNA, picoviruses use linear, monomeric DNA substrates with a terminal protein (TP) bound to each 5’ end (*28, 61*). The terminal protein gets packaged along with the DNA and is ejected with the DNA during infection. Packaging is carried out by a “packaging motor” ATPase (gp8 in Andhra). The TP is also responsible for recruitment of the DNA polymerase and initiation of replication (*28, 31*). In ϕ29, the gene encoding TP (ORF3) immediately precedes the DNA polymerase gene (Fig. 1). Vybiral et al (*30*) suggested that the P68 terminal protein was gp11 (equivalent to Andhra gp10), but gp10 encodes a lysin (*32*) that we have identified as part of the tail tip (Fig. 5). Based on structural analysis, the most likely candidate for the Andhra TP is gp7 (Fig. 5H). Density observed inside the tail knob thus most likely corresponds to this protein (Fig. 5G).

Since WTA is often the primary receptor for staphylococcal phages, the type of WTA present on the surface of the host is a key host range determinant (*62–64*). Most strains of *S. aureus* has WTA based on a phospho-glycerol (GroP) polymer backbone, whereas *S. epidermidis* and other coagulase-negative staphylococci have WTA based on a phospho-ribitol (RboP) backbone (*35, 65, 66*). In both cases, this backbone is derivatized by various sugars and amino acids. We previously isolated several picoviruses with specificity for either *S. aureus* or *S. epidermidis* (*32, 33*). We analyzed the sequences of RBPs from several of these, and show that they fall into two distinct groups, commensurate with the type of WTA of their hosts: those that infect *S. aureus* have RBPs similar to that of P68, while those that infect *S. epidermidis* have RBPs similar to that of Andhra (Table 3). The P68 RBP is closely related to that of the *S. aureus* infecting siphovirus 80α (*42*), suggesting that the RBP structure in general follows the host rather than the lineage of the phage. The slight homology between the two groups corresponds to the conserved N-terminal sequence, where the proteins interact with the more conserved portal and stem proteins (Fig. S5E,F).

**Table 3.** Sequence identity between the RBPs of various picoviruses. Percentage identity was calculated in Clustal Omega. The two clusters of viruses that infect *S. epidermidis* and *S. aureus* are highlighted in red and blue, respectively. GenBank IDs are: Andhra, KY442063.1; Pontiff, MH972262.1; Jbug18, MH972263.1; Pabna, MH972260.1; 44AHJD, AF513032.1; P68, AF513033.1.

|  | Andhra | Pontiff | JBug18 | Pabna | 44AHJD | P68 |
| --- | --- | --- | --- | --- | --- | --- |
| Andhra | 100.00 |  |  |  |  |  |
| Pontiff | 97.37 | 100.00 |  |  |  |  |
| JBug18 | 97.04 | 99.34 | 100.00 |  |  |  |
| Pabna | 23.45 | 23.45 | 23.24 | 100.00 |  |  |
| 44AHJD | 23.03 | 23.03 | 22.81 | 95.66 | 100.00 |  |
| P68 | 23.03 | 23.03 | 22.81 | 95.66 | 100.00 | 100.00 |

Binding to WTA is only the first step in the infection mechanism. Enzymatic activities associated with the tail are then needed to degrade the cell wall. Several tail proteins have enzymatic activities: gp6, a predicted hydrolase/lipase; gp10, which has a glycosyl hydrolase/peptidase (GyH) domain connected to a Cys-His aminohydrolase/peptidase (CHAP) domain (*32*); and RBP, with predicted homology to alginate lyases. (Andhra gp14 is also a lysin with amidase activity, but is not associated with the tail, and is assumed to act from within during cell lysis (*32*).) It is likely that conformational changes occurring upon the initial binding of the RBPs to WTA release the lytic activities associated with gp6 and gp10 to break down the cell wall. No equivalent of gp6 exists in P68, suggesting that the associated enzymatic activity may be specific to *S. epidermidis*. The role of gp1 in the tail tip is not clear, but is most likely also involved in penetration of the cell wall. It is not known if additional, specific interactions between the tail and membrane-associated proteins in the host cell are required. Penetration of the membrane might cause the tail tip to fall apart, allowing the α-helices of gp10 to disengage from the tail knob, leading to a conformational change in the internal α-helices in the knob, similar to that described for ϕ29 (*48*). Opening up of the tail tube would then allow the terminal protein to escape, pulling the DNA along with it, similar to the process proposed for the tape measure protein of phage 80α (*42*).

Most phages currently in use for phage therapy are large, lytic myoviruses that have genomes of >150,000 base pairs that encode hundreds of genes, most with unclear functions (*23*). Furthermore, phage therapy typically employs cocktails of multiple phages. In contrast, picoviruses are attractive candidates for therapeutic use due to their small and well-defined genomes, lack of virulence factor genes, strictly lytic life cycle and unique DNA packaging strategy that precludes generalized transduction. However, staphylococcal picoviruses are typically restricted to a narrow range of hosts. This narrow host range is a double-edged sword: on one hand, it allows the phages to target only the specific pathogen causing the disease; on the other hand, it means that the phage may need to be tailored to the patient in each specific case. Genetic manipulation of lytic picoviruses like P68 and Andhra has recently been facilitated by CRISPR technology (*67*), making possible the production of custom-made phages tailored to a specific purpose. With the present study, we have gained a better understanding of the structures and functions of the Andhra gene products and the determinants of host specificity, paving the way for a more rational design of custom phages for therapeutic applications.

## Materials and Methods

### Purification and imaging of phage Andhra

Andhra was originally isolated from raw sewage and purified as previously described (*32*), yielding a lysate (≈1x 10^9^ pfu/mL) of which 150 µL were combined with 300 µL of an overnight culture of *S. epidermidis* RP62a, 7 mL semisolid “sloppy” agar (2.3 g brain heart infusion and 0.75 g agar in 250 mL water) and 7 µL of 5 M CaCl_2_ at 55°C. The mixture was plated onto tryptic soy agar with 5 mM CaCl_2_ and allowed to solidify before incubating at 37°C overnight. Complete lysis occurred, and the sloppy agar layer containing phage was collected and combined with 20 mL tryptic soy broth per plate. Samples were then vortexed and centrifuged at 4,400 g for 5 min. The supernatant was transferred to a new tube and centrifuged at 4,400 g for 5 min. After the second centrifugation, all lysates were combined and filtered via vacuum filtration. The clarified lysates were made up to 10% PEG 8,000 and 1.0 M NaCl and incubated overnight at 4 °C. The solutions were centrifuged at 11,000 x g, 45 min, and the supernatants discarded. The pellet containing Andhra was resuspended in 2 mL of phage buffer (50 mM Tris-HCl pH 7.8, 100 mM NaCl, 1 mM MgSO_4_, 4 mM CaCl_2_) by gentle shaking. DNase I (10 mg/mL) and RNase A (2 mg/mL) were added to the pellet before incubating at 37 °C for 30 min with agitation. The sample was collected and layered onto a CsCl step gradient with densities of 1.3, 1.5 and 1.7 g/mL and centrifuged in a SW41 rotor at 140,000 x g for 4 hrs at 12 °C. Visible bands were removed from the tubes and centrifuged separately in a Ti70 rotor at 185,000 x g, 4 °C for 1 hr. Supernatant was removed, and pellets were resuspended in 100 µL phage buffer at 4 °C overnight with gentle shaking. The concentrated phage was then dialyzed in phage dialysis buffer (20 mM Tris-HCl pH 7.8, 50 mM NaCl, 1 mM MgSO_4_ and 4 mM CaCl_2_).

### Mass spectrometry

Andhra in phage buffer was prepared in Novex NuPage LDS sample buffer (Invitrogen), separated on a Novex NuPage 10% Bis-Tris polyacrylamide gel (Invitrogen) and stained overnight with Novex Colloidal Blue (Invitrogen). One entire lane was divided into four fractions, which were equilibrated in 100 mM ammonium bicarbonate and digested overnight with 20 µg/mL trypsin (Trypsin Gold, Mass Spectrometry grade; Promega) or 20 µg/mL chymotrypsin prior to LC-MS analysis. The peptide digests were separated by HPLC on a C-18 reverse phase column using a 0–85% acetonitrile gradient with 0.1% formic acid, running in-line with a Thermo Q Exactive HFx mass spectrometer. The mass data was searched with SEQUEST using a database containing the predicted Andhra ORFs (GenBank ID KY442063.1) and the *S. epidermidis* RP62a sequence (GenBank ID CP000029). The list of peptides was filtered using Scaffold 5.1.2 (Protein Sciences, Portland OR) with peptide and protein thresholds of 80% and 95%, respectively.

### Cryo-EM sample preparation and data collection

Purified Andhra virions in phage dialysis buffer were applied to nickel Quantifoil R2/2 grids and plunge-frozen using an FEI Vitrobot Mark IV at 100% humidity with 5 sec blot time and blotting force 5. Grids were screened and a preliminary data set consisting of 110 movies was collected using SerialEM on an FEI Tecnai F20 microscope operated at 200 kV, equipped with a Gatan K3 detector, with 1.45 Å pixel size. Each movie consisted of 20 frames and a total electron dose of 32 e^−^/Å^2^. Frame alignment was done on the fly using the built-in SerialEM module. A data set consisting of 8,818 movies were collected with a FEI Titan Krios microscope operated at 300 kV using a Gatan K3 electron detector with a pixel size of 0.664 Å in superresolution mode, with nominal defocus range 1.0–3.5 µm. Each movie contained 40 frames and used a total electron dose of 39 e^−^/Å^2^ (Table 2). Frame alignment was done with MotionCor2.

### Three-dimensional reconstruction

Three-dimensional reconstruction was done predominantly using RELION-3, v.3.0.8 and v.3.1 (*36*). A data set consisting of 5,216 particles was picked semi-automatically from the F20 images and reduced to 4,348 particles after 2D classification. A C1 starting model was made *ab initio* in RELION. The final reconstruction from this data reached 10.6 Å resolution. For the Titan Krios data, a total of 282,862 particles were picked semi-automatically in RELION (Table 2). 2D classification yielded a data set of 230,714 particles with an appearance consistent with Andhra virions. The C1 reconstruction from the F20 data was used as a starting model for 3D classification and refinement. The final C1 reconstruction used a subset of 62,061 particles and reached 4.43 Å resolution. For icosahedral reconstruction of the capsid, the images were re-extracted to be centered on the capsid. A starting model was made by icosahedrally averaging the asymmetric reconstruction after removing the tail density. After 3D classification, the best 90,677 particles were subjected to reconstruction with icosahedral (I1) symmetry, yielding a final map at 3.50 Å resolution. For reconstruction of the tail, images were re-extracted with a center approximately in the middle of the tail as seen in the asymmetric reconstruction and with a larger box size to accommodate the length of the tail. As a starting model, the capsid density was manually erased from the asymmetric reconstruction using UCSF Chimera (*68*), and the tail density was centered in the map. The best 154,849 particles were used for 3D refinement with C6 symmetry, yielding a final map at 3.58 Å resolution. It was clear from this map that the distal part of the tail, including the knob and tail tip, was not well resolved, presumably due to ambiguity with the 12-fold symmetry of the tail stem. The data was therefore symmetry expanded to C12 using the relion_particle_symmetry_expand script, followed by masked 3D classification focused on only the distal tail. Reconstruction of the distal tail with C6 symmetry reached a resolution of 3.90 Å. This map still did not resolve the tail tip, so another focused, masked reconstruction was carried out by C6 symmetry expansion and reconstruction without the application of symmetry, reaching a resolution of 4.90 Å (Table 2).

### Atomic model building

For most of the Andhra proteins, model building was done *de novo* using *Coot* (*69*). In some cases, starting models were generated using I-TASSER (*70*) or using AlphaFold (*47*) from within ChimeraX (*71*). Real-space refinement was done in *Coot*, followed by refinement in Phenix (*72*). For the tail, refinement included 33 protein chains, while capsid refinement included 18 protein chains (one asymmetric unit plus surrounding subunits)(Table 2). All models were checked in MolProbity (*73*).

## Acknowledgements

We are grateful to Anna McGriff and Laura Parker at The University of Alabama at Birmingham and Dr. Nayeem Bari at University of Illinois Urbana-Champaign (UIUC) for help with sample preparation, Dr. Kyoko Kojima at the O’Neal Comprehensive Cancer Center’s Mass Spectrometry/Proteomics (MSP) Shared Resource at UAB for MS data collection and analysis, and Dr. Thomas Klose and Xueyong Xu at Purdue University’s Midwestern Center for Cryo-Electron Microscopy (MCCEM) for assistance with data collection. MCCEM was supported by The National Institutes of Health (NIH) grant U24 GM116789 to Dr. Wen Jiang at Purdue University. This work was supported by NIH grant R21 AI156636 to T.D. and A.H.-A.

## Author contributions

T.D. and A.H-A. provided the overall concept, design and supervision of the project. N.C.H., J.L.K and T.D. carried out the research. T.D. wrote the initial draft. T.D, N.C.H., J.L.K and A.H.-A. reviewed and edited the manuscript.

## Data Availability

The electron density maps for the reconstructions of the asymmetric virion, the icosahedral capsid, the C5 capsid, the C6 tail, the C6 tail knob and the C1 tail tip were deposited in the Electron Microscopy Data Bank with ID numbers EMD-AAA, EMD-BBB, EMD-CCC, EMD-DDD, EMD-EEE, and EMD-FFF, respectively. The corresponding atomic coordinates were submitted to the RCSB Protein Data Bank with ID numbers aaa, bbb, ccc, ddd, eee and fff. *[NOTE: At the time of manuscript submission, deposition is still in progress and ID numbers are not yet available]*

