## Supplementary figures for "Structure and host specificity of *Staphylococcus epidermidis* bacteriophage Andhra"

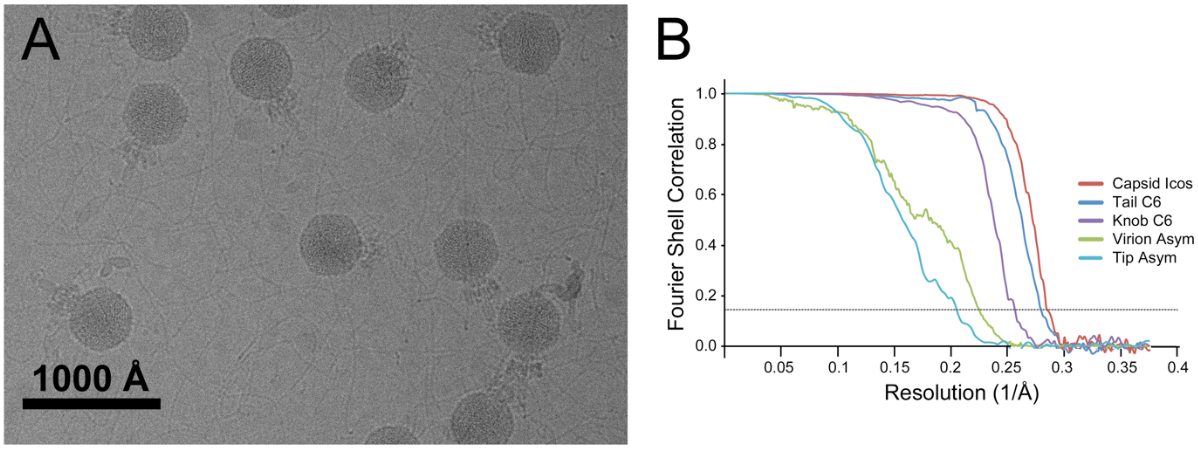


**Fig. S1. Imaging and reconstruction of phage Andhra. (A)** Cryo-electron micrograph of Andhra virions. Scale bar, 1000Å. **(B)** Fourier shell correlation curves between two half-maps for the asymmetric reconstruction of the whole virion (green), C6 reconstruction of the tail (dark blue), icosahedral reconstruction of the capsid (red), C6 reconstruction of the knob (purple), and the asymmetric reconstruction of the tail tip (light blue). The 0.143 cutoff level is shown as a dotted line.

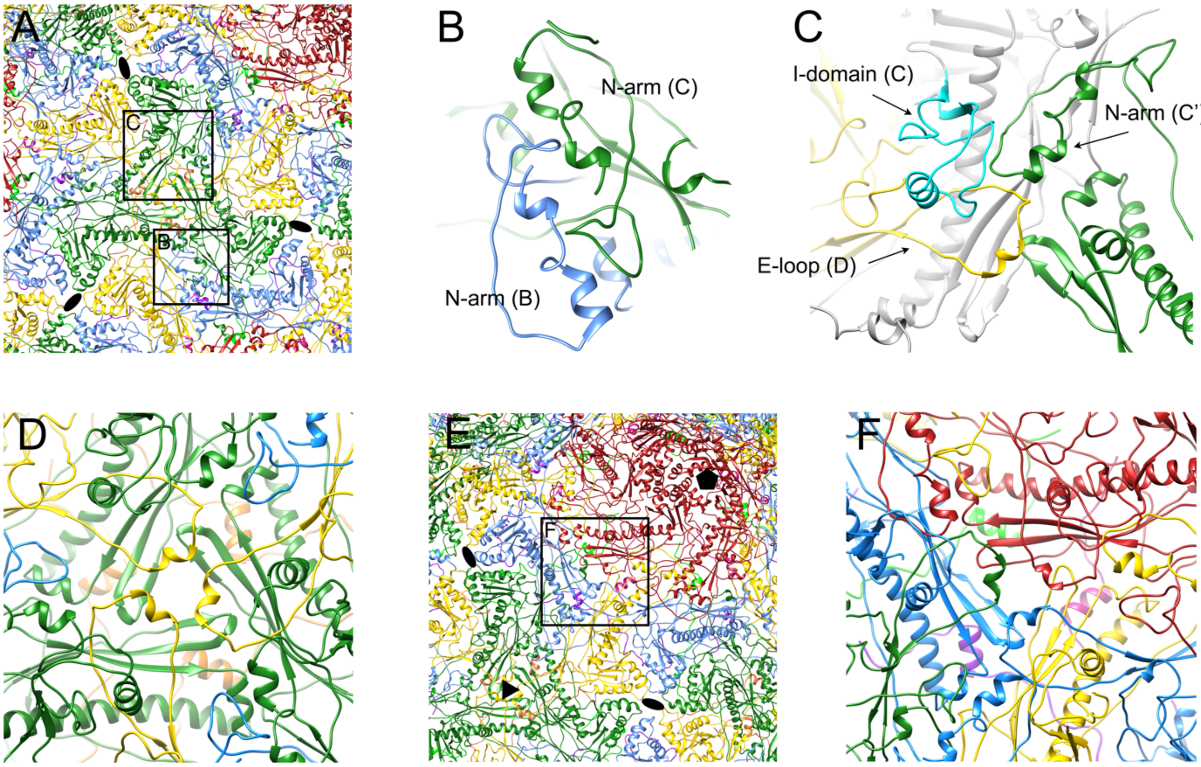


**Fig. S2. Capsid protein interactions. (A)** Capsid model, viewed down the icosahedral threefold axis between three hexamers. CP and CLP subunits are colored as in Fig. 3 (CP: A, red; B, blue; C, green; D, yellow. CLP: E, magenta; F, purple; G, light green; D, orange). Twofold axes are labeled with ellipses. The squares correspond to the views in (B) and (C). **(B)** Detail of interaction between the N-arms of a B (blue) and C (green) subunit. **(C)** Detail of the interaction between the I-domain of a C subunit (cyan, the rest of the subunit in gray), the N-arm of an adjacent C subunit (C'; green), and the E-loop of a D subunit (yellow). **(D)** Interaction between the P-domains of three C subunits (green) and the E-loops of three D subunits (yellow) around the icosahedral threefold axis. **(E)** Capsid model, viewed down the quasi-threefold axis between one pentamer and two hexamers. Twofold, threefold and fivefold axes are labeled with ellipses, triangle and pentagon, respectively. The square corresponds to the view in (F). **(F)** Closeup view of the interaction between the P domains of the A, B and D subunits and the E-loops of the A, B and C subunits around the quasi-threefold axis.

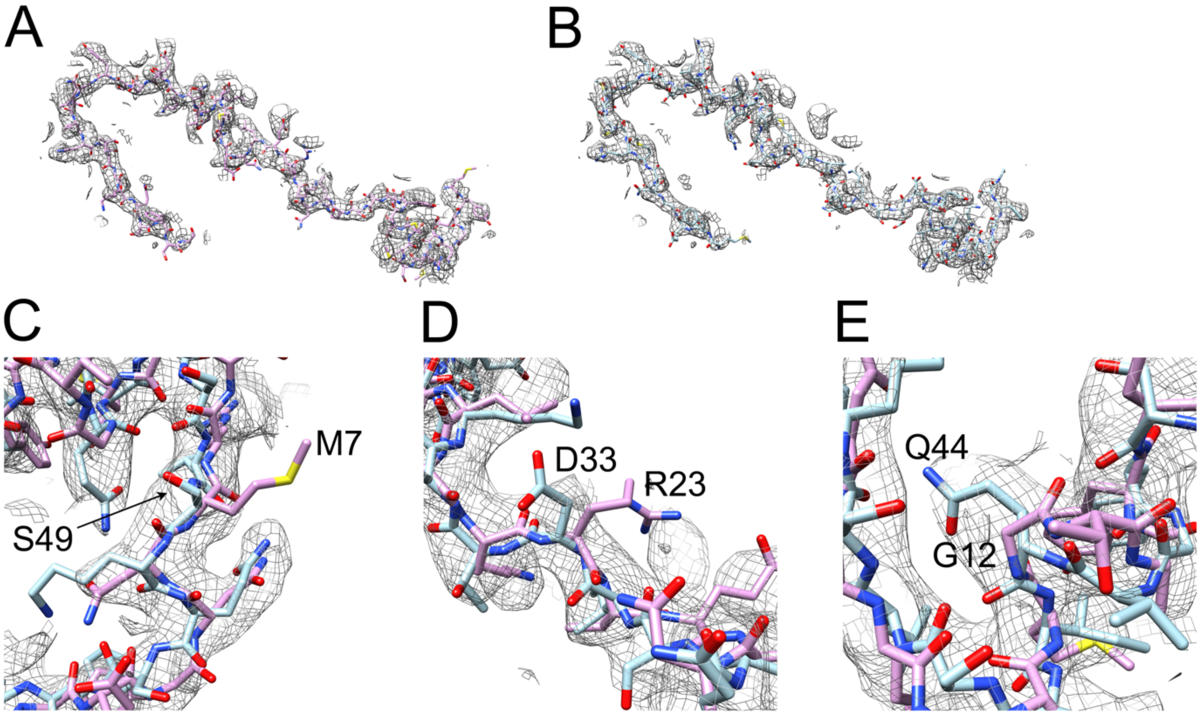


**Fig. S3. The capsid lining protein in P68. (A)** P68 gp21 as submitted (PDB ID: 6IB1) in the corresponding P68 density (EMD-4442). The protein is shown in stick representation with element colors, with the carbon backbone in pink. **(B)** P68 gp21 modeled into the P68 density in the opposite direction, corresponding to the orientation in Andhra. The backbone is shown in light blue. **(C-E)** Details of the P68 density with the submitted (pink backbone) and re-modeled (blue backbone) gp21 models. Pertinent residues in both models are highlighted.

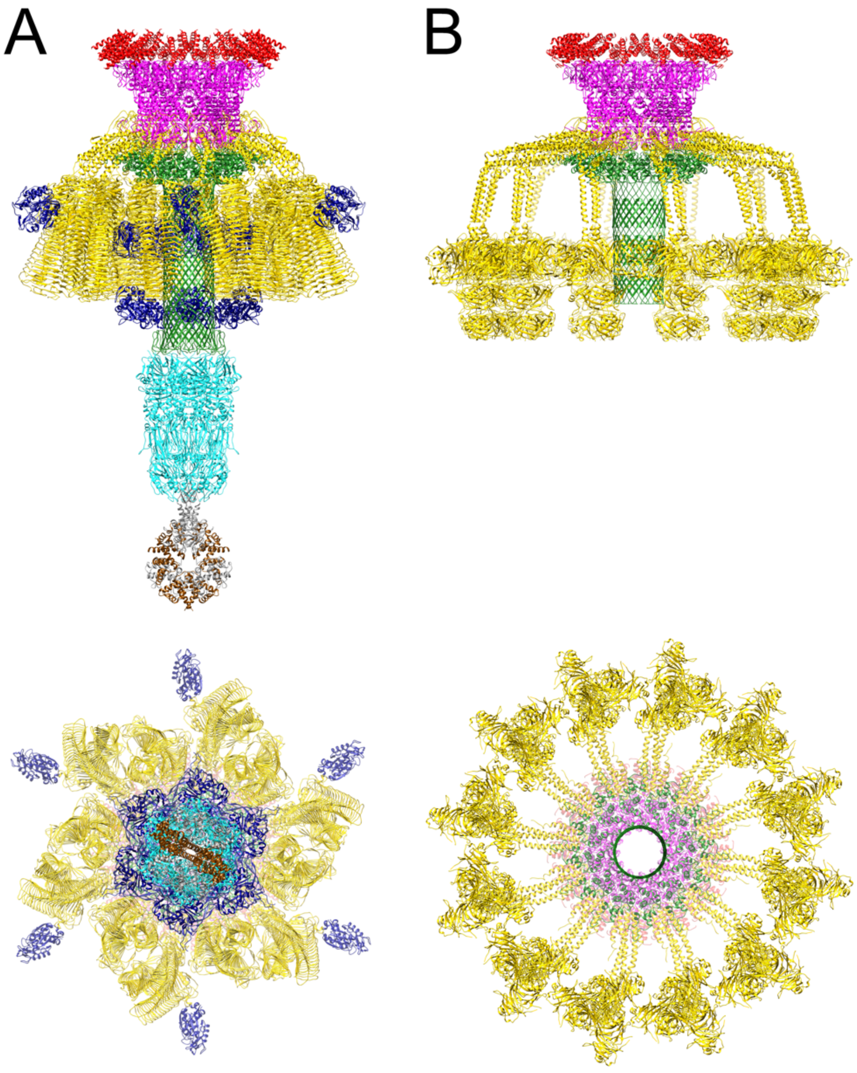


**Fig. S4. Comparison of Andhra and P68 structures.** Ribbon diagram of the composite model of Andhra **(A)** compared to P68 **(B)**, viewed form the side (top panel) and from the tip of the tail (bottom panel). Colored as in Figs 1 and 4.

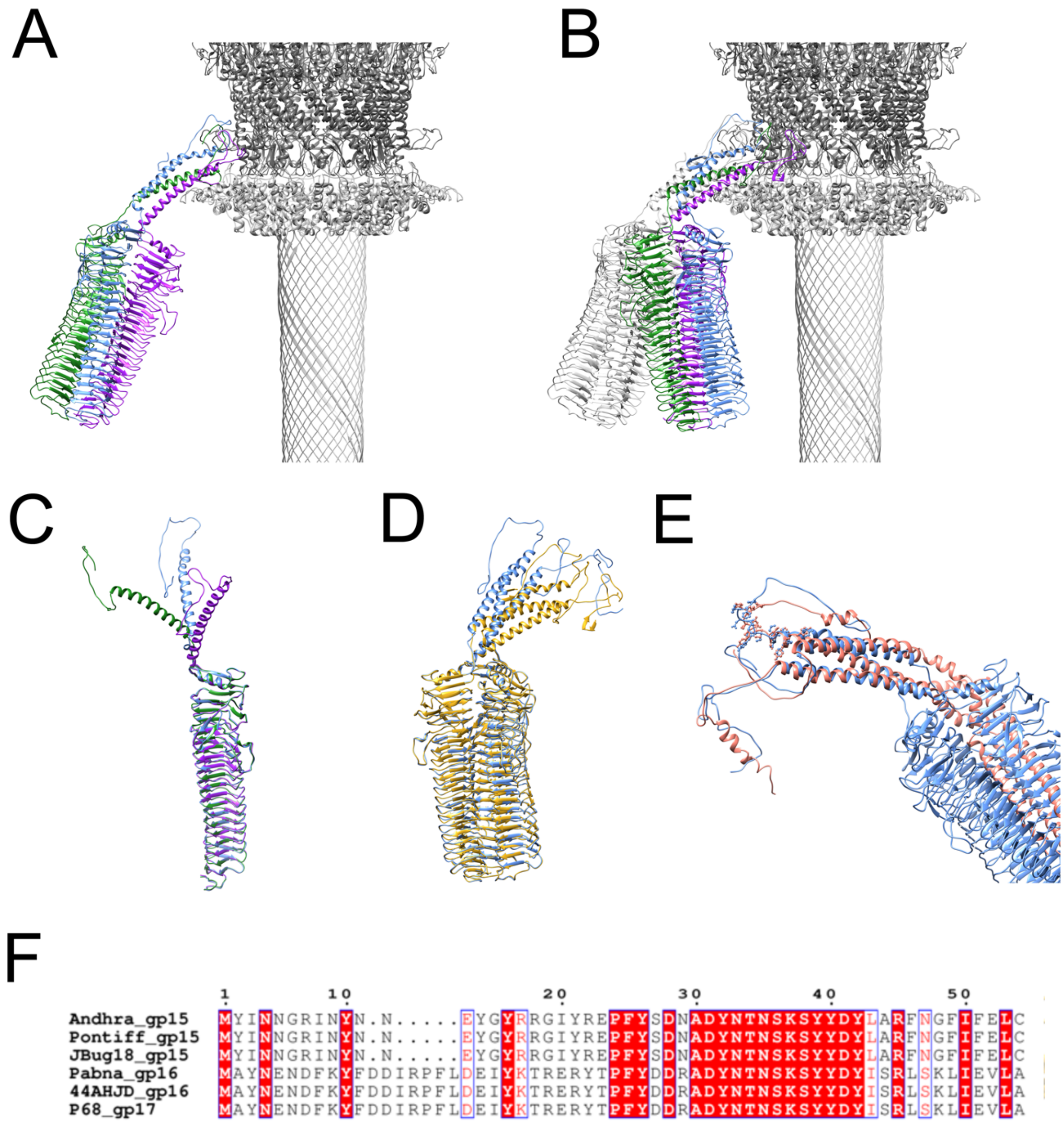


**Fig. S5. The receptor binding protein. (A)** Ribbon diagram showing the interaction of the distal RBP trimer (subunits colored blue, green and purple) with the stem and portal proteins (light and dark gray, respectively). **(B)** Interaction of the proximal RBP trimer with the stem and portal, colored as in (A). The distal RBP trimer is shown in gray. **(C)** Superposition of the three protomers in the distal RBP trimer, showing the distinct path of the N-terminal arms, colored as in (A). **(D)** Superposition of the distal (gold) and proximal (blue) RBP trimers. **(E)** Superposition of the N-terminal domains of Andhra RBP (blue) with the P68 RBP (salmon). The stretch of conserved residues in one of the subunits in both Andhra and P68 is shown in ball-and-stick representation.

**(F)** Alignment of the N-terminal sequences of the RBPs from Andhra, Pontiff, JBug18, Pabna, 44AHJD and P68. Red boxes indicate 100% sequence identity.
